# H3K4me3 is a post-transcriptional histone mark

**DOI:** 10.1101/2025.07.03.662929

**Authors:** Komal Paresh Walvekar, Sabarinadh Chilaka

**Affiliations:** Division of Applied Biology, CSIR-Indian Institute of Chemical Technology, Hyderabad-500007, Telangana, India; Academy of Scientific and Innovative Research (AcSIR), Ghaziabad, Uttar Pradesh-201 002, India; DBT-Ramalingaswami Re-entry Fellow, Department of Biotechnology (DBT), Regional Centre for Biotechnology, Faridabad, Haryana 121 001, India

**Keywords:** H3K4me3, transcriptional regulation, temporal dynamics, inducible gene model, and THP-1 monocytes

## Abstract

Histone H3 Lysine 4 trimethylation (H3K4me3) is widely recognized as a hallmark of actively transcribed gene promoters, yet its precise role in transcriptional regulation remains unresolved^1–4^. Here we present the first direct evidence that H3K4me3 is not a cause but a consequence of transcriptional activation, functioning as a downstream epigenetic response. Using inducible gene models to study temporally resolved transcriptional dynamics, we demonstrate that H3K4me3 enrichment significantly lags behind the transcriptional peaks of inflammatory genes such as *TNF-α* and *IL-1β*, appearing hours after maximal RNA synthesis. This delayed deposition contrasts sharply with the early accumulation of other active histone marks, including H3K27ac and H4K8ac, and with the recruitment of canonical transcriptional regulators such as phosphorylated RNA polymerase II, NF-κB, and chromatin modifier, histone acetyltransferase p300. Pharmacological inhibition of transcription with Actinomycin D reduces H3K4me3 enrichment, while knockdown of *MLL1* abolishes H3K4me3 without affecting gene activation, indicating that H3K4me3 deposition is transcription-dependent but dispensable for transcriptional initiation. Furthermore, analysis of the constitutively active *MYC* gene suggests that post-transcriptional H3K4me3 deposition may be a general feature of active transcription. Together, these findings redefine H3K4me3 as a post-transcriptional histone mark, revealing an unexpected layer of epigenetic regulation with broad implications for gene expression control.

## Introduction

Histone post-translational modifications play a pivotal role in regulating chromatin structure and function, profoundly influencing the gene expression programs that govern embryonic development, cell lineage specification, and the onset of pathological conditions^4–8^. Among these, H3K4me3 has emerged as one of the extensively studied histone modification, with over 4,000 publications in PubMed reflecting its central importance in epigenetics. Although H3K4me3 has been well-studied and considered as a hallmark of active transcription, its precise role in transcriptional regulation remains poorly understood. H3K4me3 is ubiquitously found at the promoters of actively transcribed eukaryotic genes and closely associated with open chromatin regions^9–12^.

Owing to its strong association with actively transcribed promoters, several studies have implicated H3K4me3 in facilitating transcriptional initiation^13,14^. However, direct evidence supporting this function is missing. Conversely, emerging evidence challenges this paradigm, as studies across diverse biological contexts, including development and specific gene classes, have shown that transcription can initiate in the absence of detectable H3K4me3, suggesting it may not strictly required for transcriptional activation^15–20^. This underscore a striking paradox that although H3K4me3 is strongly associated with active promoters, it appears to be dispensable for transcription initiation. This discrepancy has prompted a significant gap in our understanding of H3K4me3’s role in transcription and raises a fundamental question that does H3K4me3 actively drive transcription, or is it merely a downstream consequence of transcriptional activity^1–4,21^? Resolving this uncertainty has broad implications, from elucidating fundamental mechanisms of gene activation to informing therapeutic strategies, particularly in diseases such as cancer where H3K4me3 is frequently dysregulated.

Here we report an integrated temporal dynamic characterization of H3K4me3 during transcriptional initiation using inducible genes as a model. We uncovered an unprecedented deposition pattern for H3K4me3, demonstrating that it is loaded post-transcriptionally rather than pre-transcriptionally. Specifically, H3K4me3 deposition at the promoters of induced genes occurs after the transcriptional peak and is dependent on RNA synthesis. These findings provide the first direct evidence that challenges the traditional view of H3K4me3 as a marker of active transcription initiation, thereby advancing a novel conceptual framework for its involvement in the epigenetic regulation of gene expression.

## Results

### Inducible transcription model to investigate H3K4 methylation dynamics

To better understand the dynamic regulation of H3K4me3 during transcription initiation, we utilized an inducible gene system as a precise and innovative model for studying transcriptional activation. We propose that, unlike constitutively active gene models, inducible gene models provide precise control over gene off-to-on transitions, making them ideal for studying the dynamic regulation of histone post-translational modifications. Here, we employed the well-characterized human inducible inflammatory genes tumor necrosis factor-alpha (*TNF-α*) and interleukin-1 beta (*IL-1β*) to investigate histone dynamics associated with transcriptional activation. These genes undergo robust transcriptional induction in immune cells, such as monocytes and macrophages, upon stimulation with inflammatory agents like bacterial lipopolysaccharide (LPS)^22–25^. Human monocytic THP-1 cells were treated with LPS for various time points (0, 0.5, 2, 6, 12, and 24 h), after which they were harvested for parallel analysis of transcriptional activation and profiling of histone modifications at the corresponding promoters using quantitative RT-PCR (qRT-PCR) and chromatin immunoprecipitation (ChIP) assays, respectively. (Fig. 1a). qRT-PCR analysis showed a significant upregulation of *TNF-α* and *IL-1β* mRNA levels compared to 0 h, with peak expression observed at 2 h following LPS stimulation (Fig. 1b and Supplementary Fig. 1a). This transcriptional upregulation was transient, as mRNA levels began to decline noticeably by 6 h and continued to decrease at 12 and 24 h. The observed rapid and transient expression patterns of *TNF-α* and *IL-1β* is consistent with their well-established temporal response to LPS stimulation^22–24^.

**Figure 1.**
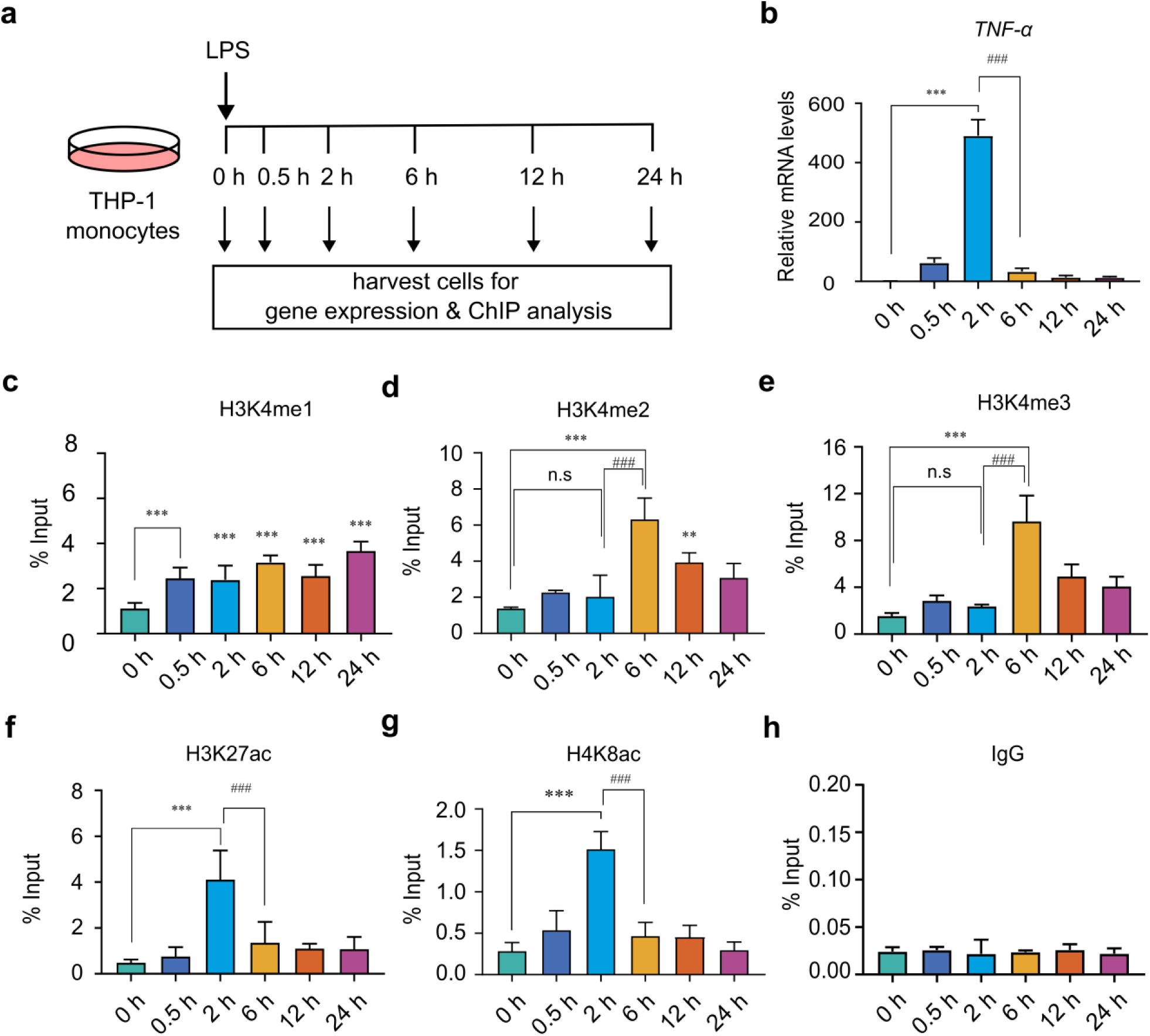
Inducible gene activation model reveals temporal decoupling of H3K4me3 deposition and transcription. (**a**), Schematic of the experimental model used to study the temporal dynamics of histone post-translational modifications during LPS-induced transcriptional activation. Cells were stimulated with LPS (500 ng/ml) and harvested at 0, 0.5, 2, 6, 12, and 24 h post-stimulation for transcriptional and chromatin immunoprecipitation (ChIP) analyses. (**b**), qPCR analysis of *TNF-α* mRNA levels following LPS stimulation, used as a readout of transcriptional activation. (**c–h**), ChIP enrichment analysis was performed using antibodies against (**c**) H3K4me1, (**d**) H3K4me2, (**e**) H3K4me3, (**f**) H3K27ac, (**g**) H4K8ac, and (**h**) IgG control. Immunoprecipitated DNA was analysed by qPCR using primers specific to the *TNF-α* promoter. Statistical analysis was performed using one-way ANOVA followed by Dunnett’s multiple comparisons test. Data are presented as mean ± SD (n=3) and SEM (n=4 for b) from independent biological replicates. Statistical significance is indicated as \*\*\**p* < 0.001, \*\**p* < 0.01, and \**p* < 0.05, relative to the unstimulated control, ###P < 0.001, ##P < 0.01, #P < 0.05 compared to LPS 2 h stimulated cells and ns as non-significant compared to unstimulated control.

### Histone dynamics analysis revealed that H3K4me3 is deposited post-transcriptionally

H3K4 methylation, comprising H3K4me1, H3K4me2, and H3K4me3, is a well characterized histone modification closely linked to active gene transcription. H3K4me1 is commonly associated with active enhancers, H3K4me2 is predominantly enriched near the 5′ end of actively transcribed genes, and H3K4me3 typically marks the promoters of actively transcribed genes^12^. To elucidate the dynamic regulation of H3K4 methylation in relation to transcription initiation, we conducted chromatin enrichment analysis of H3K4me1, H3K4me2, and H3K4me3 at the *TNF-α* and *IL-1β* promoters during transcriptional activation (Fig. 1a). ChIP-qPCR analysis revealed a significant increase in H3K4me1 at the *TNF-α* promoter as early as 0.5 hours following LPS induction, with elevated levels persisting at later time points (Fig. 1c). Surprisingly, H3K4me2 and H3K4me3 did not exhibit enrichment at the *TNF-α* promoter during the peak of transcription observed 2 h after induction (Fig. 1d-e). Notably, strong and highly significant enrichment of both marks was observed at 6 h post-induction, despite a marked reduction in transcriptional activity (Fig. 1d–e). To further support these findings, ChIP enrichment analyses were performed for additional histone marks associated with active transcription, including H3K27ac and H4K8ac^26,27^. ChIP-qPCR results showed a significant increase in H3K27ac and H4K8ac marks at 2 h, coinciding with peak *TNF-α* mRNA expression (Fig. 1f–g). This aligns with the well-established role of histone acetylation in promoter activation during transcription and further validates our transcriptional model. Consistent with the H3K4 methylation dynamics observed at the *TNF-α* promoter, *IL-1β* transcriptional activation exhibited a similar pattern of histone methylation changes (Supplementary Fig. 1b–g), further substantiating these findings. Overall, these results demonstrate that H3K4me2 and H3K4me3 deposition did not correlate with the *TNF-α* and *IL-1β* transcriptional peak at 2 h, but showed a delayed enrichment post-transcriptionally at 6 h, irrespective of gene activity.

### Transcriptional activators display chromatin binding patterns distinct from H3K4me3

To further support the observation that H3K4me3 enrichment represents a post-transcriptional event uncoupled from transcriptional activity, we analysed whether the 6 h time point corresponds to a transcriptionally inactive state by examining the temporal dynamics of key transcriptional markers. Phosphorylation of the C-terminal domain (CTD) of RNA polymerase II (Pol II) at serine 5 (Ser5P) and serine 2 (Ser2P) serves as a well-established indicator of transcriptional activity^28^. Specifically, Ser5P is associated with transcription initiation, while Ser2P marks the elongation phase. Chromatin enrichment analysis of Pol II Ser5P at the *TNF-α* and *IL-1β* promoters revealed a significant increase at the 2 h time point compared to the 0 h control (Fig. 2a and Supplementary Fig. 2a), consistent with active transcription initiation. However, Ser5P levels were significantly reduced at 6 h compared to 2 h, indicating a rapid downregulation of transcriptional initiation activity (Fig. 2a and Supplementary Fig. 2a). Similarly, enrichment analysis of Pol II Ser2P at the *TNF-α* gene body (exon 4) showed a significant increase at 2 h, coinciding with the peak of transcriptional activity, followed by a significant decrease at 6 h compared to the 0 h control (Fig. 2b and Fig. 1b), suggesting a rapid decline in transcriptional elongation. At later time points (12 and 24 h), both Ser5P and Ser2P levels remained low, indicating sustained transcriptional inactivity. To confirm assay specificity, we assessed Ser5P and Ser2P levels at the *GAPDH* promoter, a constitutively expressed gene commonly used as a positive control. As expected, both phosphorylation marks were consistently maintained at this locus (Supplementary Fig. 3a–b), validating assay specificity and supporting the reliability of the transcriptional dynamics observed at the target loci. Taken together, the comparison of RNA Pol II Ser5P and Ser2P enrichment dynamics with H3K4me3 supports the conclusion that H3K4me3 loading at promoters occurs through a delayed post-transcriptional enrichment mechanism.

**Figure 2.**
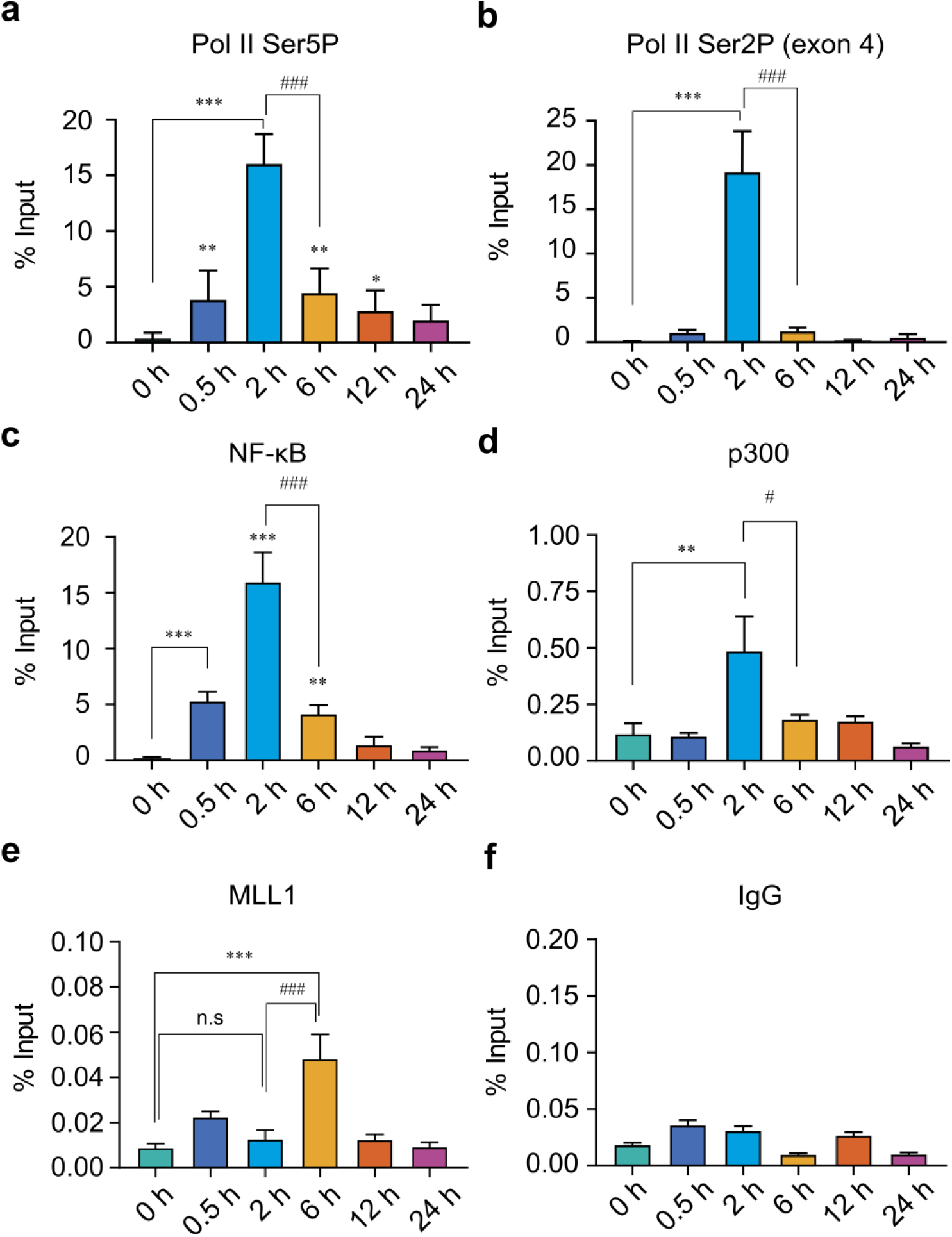
ChIP enrichment analysis of transcriptional regulators reveals temporal divergence from H3K4me3 dynamics during LPS-induced gene activation. (**a–f**), THP-1 cells were stimulated with LPS (500 ng/ml) and harvested at 0, 0.5, 2, 6, 12, and 24 hours to assess temporal occupancy of key transcriptional regulators at the *TNF-α* locus. ChIP enrichment analysis was performed using antibodies against (a) RNA polymerase II Ser5-phosphorylated (Pol II Ser5P), marking transcriptional initiation; (b) RNA polymerase II Ser2-phosphorylated (Pol II Ser2P), marking elongation; (c) NF-κB (p65), a stimulus-responsive transcription factor; (d) MLL1, a histone methyltransferase responsible for H3K4me3 methylation; (e) P300, a histone acetyltransferase linked to transcriptional coactivation; and (f) IgG as a negative control. Enriched chromatin was quantified by qPCR using primers specific to the *TNF-α* promoter (a, c–e) or exon 4 (b). Statistical analysis was performed using one-way ANOVA followed by Dunnett’s multiple comparisons test. Data are presented as mean ± SD (n =4 for a, b and n=3 for c, d and f) and SEM (n=3 for e) independent biological replicates. Statistical significance is indicated as \*\*\**p* < 0.001, \*\**p* < 0.01, and \**p* < 0.05, compared to unstimulated control, ###P < 0.001, ##P < 0.01, #P < 0.05 compared to LPS 2 h stimulated cells.

Inflammation-induced transcriptional activation of *TNF-α* and *IL-1β* is primarily mediated by NF-κB, a key regulator of inflammatory gene expression^29^. In response to inflammatory stimuli, NF-κB translocate into the nucleus and activates transcription by binding to gene promoters and recruiting co-activators such as the histone acetyltransferase p300. To elucidate the temporal coordination between NF-κB mediated transcriptional regulation and H3K4me3 deposition, we performed ChIP assays to assess NF-κB and p300 occupancy at the *TNF-α* and *IL-1β* promoters following LPS stimulation. NF-κB enrichment at the *TNF-α* promoter was detectable as early as 0.5 h post-stimulation and increased markedly by 2 h, coinciding with elevated transcriptional activity (Fig. 2c). Notably, NF-κB occupancy declined significantly by 6 h compared to 2 h, correlating with reduced transcriptional activity, indicative of its transient promoter engagement. A similar pattern of NF-κB binding was observed at the *IL-1β* promoter (Supplementary Fig. 2b), reinforcing the rapid and dynamic nature of NF-κB recruitment correlating with transcriptional induction. Furthermore, p300 occupancy exhibited a similar recruitment pattern, with peak enrichment at 2 h post-stimulation (Fig. 2d and Supplementary Fig. 2c), aligning with both NF-κB binding and peak transcriptional activity. Together, these findings demonstrate that NF-κB and p300 are recruited during the early phase of *TNF-α* and *IL-1β* transcriptional activation, in contrast to the delayed deposition of H3K4me3. This temporal separation supports our finding that H3K4me3 functions as a downstream marker of active transcription rather than a prerequisite for its initiation and may be deposited through a mechanism independent of these transcription factors.

### MLL1 recruitment correlates with post-transcriptional H3K4me3 enrichment

To independently confirm the timing of H3K4me3 deposition at the *TNF-α* and *IL-1β* promoters during transcriptional activation, we examined the recruitment of MLL1, a histone methyltransferase that catalyses H3K4 trimethylation at these loci. MLL1 was shown to promote inflammatory gene expression by H3K4me3 deposition downstream of NF-κB signalling, through its recruitment to NF-κB-bound promoters in monocytes and macrophages^30–32^. ChIP analysis showed no significant increase in MLL1 occupancy at the *TNF-α* and *IL-1β* promoters at 0.5 or 2 hours after stimulation, as MLL1 occupancy levels remained comparable to those in unstimulated controls (Fig. 2e and Supplementary Fig. 2d). Notably, by 6 h post-induction, a significant increase in MLL1 occupancy was observed, which coincided with a pronounced enrichment of H3K4me3 (Fig. 2e, Supplementary Fig. 2d, and Fig. 1e). However, at later time points (12 and 24 h), MLL1 binding decreased, paralleling the decline in H3K4me3 levels. As a positive control, we analysed MLL1 occupancy at the constitutively active GAPDH promoter, where MLL1 levels remained unchanged across all time points (Supplementary Fig. 3c–d). Collectively, these data reveal a temporal correlation between MLL1 recruitment and H3K4me3 enrichment, supporting the conclusion that H3K4me3 is deposited post-transcriptionally. The delayed recruitment of MLL1 suggests a regulatory mechanism in which MLL1 is engaged after transcription initiation, potentially promoting H3K4me3-mediated post-transcriptional chromatin remodelling.

### Transcription is a prerequisite for H3K4me3 deposition

Although H3K4me3 is enriched at active promoters, our data suggest that its accumulation occurs post-transcriptionally. We therefore hypothesized that its deposition may nonetheless depend on transcriptional activity. To evaluate this, we employed Actinomycin D (ActD), a potent transcriptional inhibitor that intercalates into DNA and disrupts RNA polymerase function^33^. THP-1 cells were stimulated with LPS in the presence or absence of Actinomycin D, and analysed at 2 and 6 h post-stimulation for *TNF-α* and *IL-1β* transcription induction and H3K4me3 enrichment at their respective promoters. LPS stimulation of naïve THP-1 cells (Naïve-LPS) induced *TNF-α* mRNA expression at 2 h, consistent with a transient inflammatory response, which diminished by 6 h (Fig. 3a). In contrast, Actinomycin D-treated cells stimulated with LPS (ActD-LPS), exhibited a marked reduction in *TNF-α* mRNA levels at 2 h compared to Naïve-LPS cells, confirming effective transcriptional inhibition (Fig. 3a). By 6 h, *TNF-α* mRNA levels in ActD-LPS cells remained low, similar to those in Naïve-LPS cells. Consistent with the H3K4me3 dynamics observed above, H3K4me3 enrichment at the *TNF-α* promoter in Naïve-LPS cells was markedly increased at 6 h, but not at 2 h post-stimulation (Fig. 3b). Interestingly, ActD-LPS cells exhibited significantly reduced H3K4me3 enrichment at 6 h compared to Naïve-LPS cells, suggesting that transcriptional inhibition impairs H3K4me3 deposition (Fig. 3b). A similar effect was observed at the *IL-1β* promoter, where transcriptional blockade at 2 h impaired H3K4me3 enrichment by 6 h (Supplementary Fig. 4a–b). Together, these findings indicate that active transcription is necessary for the deposition of H3K4me3 at gene promoters.

**Figure 3.**
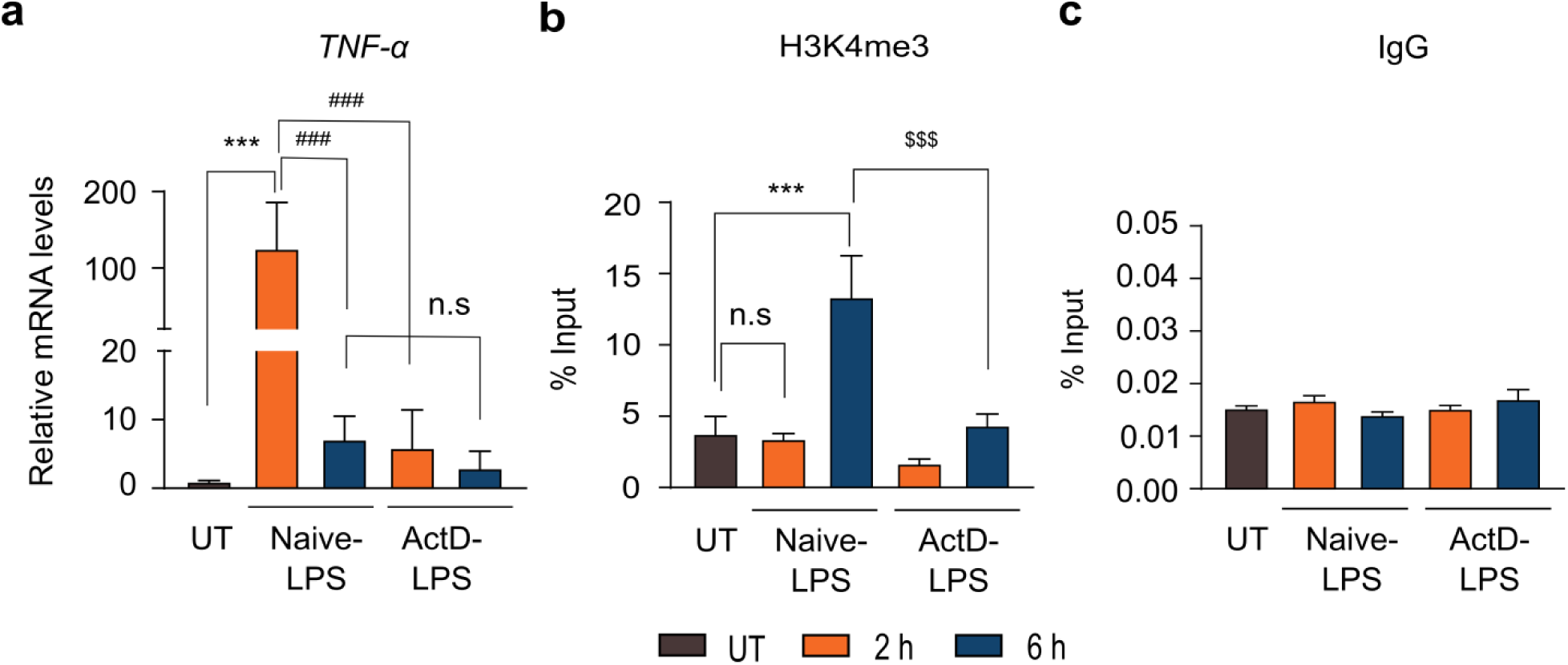
Post-transcriptional H3K4me3 loading requires active transcription. To determine whether active transcription is necessary for the post-transcriptional deposition of H3K4me3 at gene promoter, THP-1 monocytic cells were pre-treated with the transcriptional inhibitor Actinomycin D (1 µM) for 30 min prior to stimulation with LPS (500 ng/mL), and cells were harvested at 0 h, 2 h, and 6 h post-stimulation for analysis of gene expression and histone modifications. (**a**) Quantitative RT–PCR analysis of *TNF-α* mRNA levels in control and Actinomycin D-treated THP-1 cells following LPS stimulation. (**b**) ChIP analysis using antibodies against H3K4me3 to assess enrichment at the *TNF-α* promoter. (**c**) ChIP with normal IgG was included as a negative control. Statistical analysis was performed using one-way ANOVA followed by Dunnett’s multiple comparisons test. Data are presented as mean ± SD (n =3 for a and c), SEM (n=4 for b) independent biological replicates. Statistical significance is denoted as P < 0.001, P < 0.01, P < 0.05 versus unstimulated control, ###P < 0.001, ##P < 0.01, #P < 0.05 compared to LPS 2 h stimulated cells, $$$P < 0.001, $$P < 0.01, $P < 0.05 compared to LPS 6 h stimulated cells and ns as non-significant compared to unstimulated control.

### H3K4me3 is dispensable for the transcriptional activation of *TNF-α*

Although H3K4me3 has been shown to be non-essential for transcription initiation¹⁵-¹⁹, the molecular basis for this observation has remained unclear. Here, we show that H3K4me3 is deposited after the peak of transcriptional activity, explaining its non-essential role in transcription initiation. To demonstrate this dispensability in the context of the current inducible gene activation model, we analysed the transcriptional activation of *TNF-α* under conditions of reduced H3K4me3 levels. We used shRNA to deplete MLL1, a histone methyltransferase recruited to NF-κB-bound promoters and implicated in inflammatory gene activation in monocytes and macrophages^30–32^. Stable THP-1 monocytic cell lines expressing shRNAs (n=2) targeting human *MLL1* were generated using the pLKO.1 lentiviral vector system. Western blot analysis confirmed efficient depletion of MLL1, accompanied by a marked global reduction in H3K4me3 levels compared to control cells transduced with the empty vector (Fig. 4a-b). Upon LPS stimulation, MLL1 knockdown cells exhibited *TNF-α* mRNA levels comparable to those of control cells at 2 h (Fig. 4c), indicating that reduced global H3K4me3 due to MLL1 knockdown does not impair *TNF-α* induction. These results support the conclusion that H3K4me3 is dispensable for the initiation of *TNF-α* transcription. Notably, MLL1 knockdown cells exhibited a progressive decline in viability and proliferation over time, suggesting that MLL1 and its associated histone mark, H3K4me3, may contribute to essential cellular functions (data not shown).

**Figure 4.**
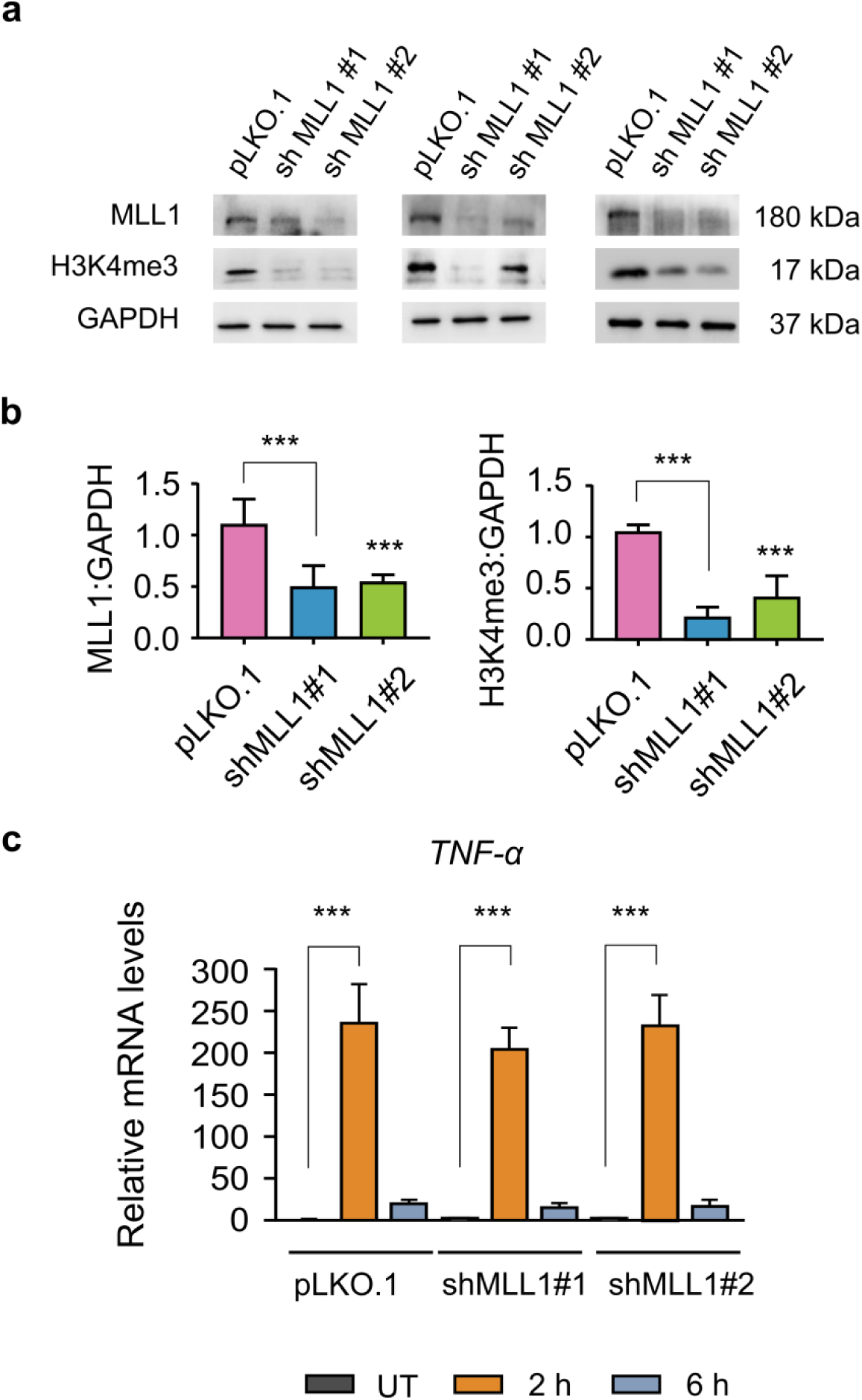
H3K4me3 depletion does not impair LPS-induced *TNF-α* mRNA expression. THP-1 cells with MLL1 knockdown were generated by lentiviral transduction using pLKO.1 vectors expressing two independent shRNAs. Transduced cells were selected with puromycin (1 µg/ml) for 7 days to establish stable cell populations. Following selection, cells were used for LPS-induced transcriptional analysis of *TNF-α.* (a), Western blot analysis of MLL1 and H3K4me3 levels in control and shMLL1 cells. GAPDH served as a loading control. (b), Densitometric quantification of immunoblot signals performed using ImageJ. (c), Quantitative RT–PCR analysis of TNF-α mRNA expression in control and shMLL1 cells following LPS stimulation (2 h and 6 h). Statistical analysis was performed using one-way ANOVA with Dunnett’s multiple comparisons test. Data are presented as mean ± SD from three independent biological replicates (n = 3). Statistical significance is indicated as **P < 0.001, *P < 0.01, and P < 0.05 compared to unstimulated control.

### Post-transcriptional H3K4me3 enrichment reflects a broad epigenetic mechanism

Given that post-transcriptional H3K4me3 deposition was observed at inducible genes, we next investigated whether this feature is also characteristic of constitutively expressed genes. However, studying delayed transcriptional effects, such as H3K4me3 deposition, at constitutively active genes is inherently challenging because continuous transcription can mask subtle temporal dynamics. To overcome this limitation, we used transcriptional termination as a model system to investigate the dynamics of H3K4me3 deposition. We hypothesise that if H3K4me3 is deposited post-transcriptionally with a delay following transcriptional activity, it should remain at the promoter for an extended period even after transcription has terminated (Fig. 5a). We selected the *MYC* gene for this analysis because it is constitutively expressed and, importantly, has a very short mRNA half-life^34^. The short half-life of *MYC* mRNA makes it a sensitive and reliable indicator of transcriptional status, as residual transcript levels following Actinomycin D treatment closely reflect the degree of ongoing transcription. To assess transcriptional dynamics and epigenetic changes, THP-1 cells were exposed to Actinomycin D for 0, 1, 2, 4, 6, or 8 h, followed by quantification of *MYC* mRNA and H3K4me3 levels at the *MYC* promoter. Following Actinomycin D treatment, *MYC* mRNA levels declined to approximately 20% by 2 h, less than 10% by 4 h, and below 5% at 6 and 8 h, compared to untreated cells at 0 h, demonstrating effective transcriptional inhibition (Fig. 5b). ChIP enrichment analysis of H3K4me3 at the *MYC* promoter revealed sustained levels up to 4 h of treatment, comparable to those observed in the untreated control (Fig. 5c). Notably, its levels were declined significantly at 6 and 8 h (Fig. 5c). The sustained H3K4me3 levels at 2 and 4 h, despite effective transcriptional shutdown, indicate a delayed deposition of H3K4me3. The uncoupling of transcription and H3K4me3 dynamics at the *MYC* promoter indicates that post-transcriptional H3K4me3 deposition may represent a broader epigenetic mechanism of transcriptional regulation.

**Figure 5.**
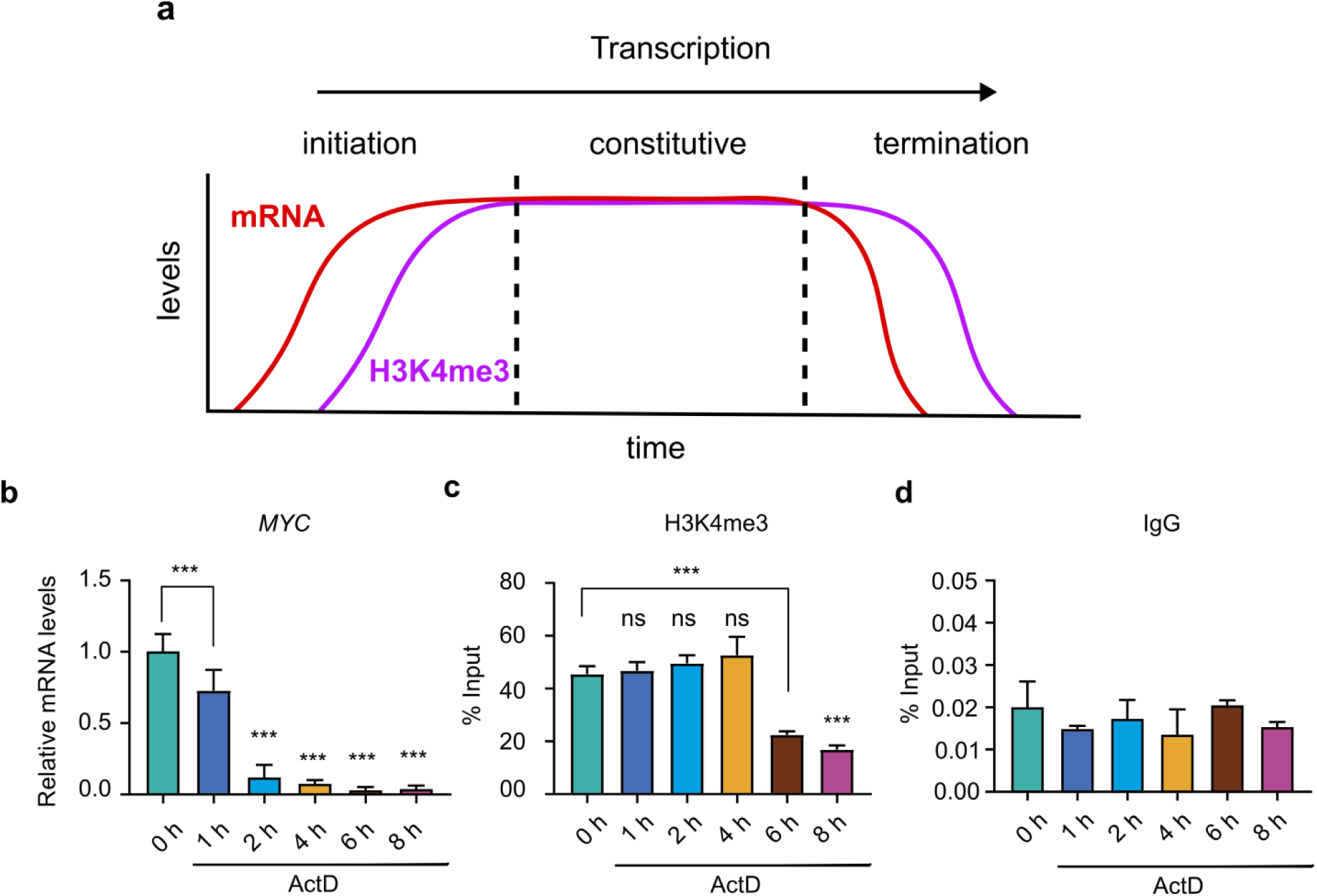
Sustained H3K4me3 enrichment at the constitutively active *MYC* promoter despite transcriptional inhibition suggests delayed deposition. (**a**) Schematic representation illustrating the hypothesized deposition of H3K4me3 at the TSS during distinct transcriptional phases, including initiation, constitutive expression, and termination. To assess whether post-transcriptional delay in H3K4me3 deposition occurs at the constitutively active *MYC* promoter, THP-1 cells were treated with Actinomycin D (1 µM) for the indicated durations (0.5, 1, 2, 6, and 8 h), and MYC transcription levels and H3K4me3 enrichment at the *MYC* promoter were analysed. (b), Quantitative RT–PCR analysis of *MYC* mRNA levels in THP-1 cells treated with Actinomycin D (1 µM) for the indicated durations (0.5, 1, 2, 6, and 8 h), demonstrating effective transcriptional inhibition. (c), ChIP assays were performed using antibodies against H3K4me3 to assess its enrichment at the *MYC* promoter under transcriptionally repressed conditions. (d), ChIP with isotype-matched IgG was included as a negative control. Statistical analysis was performed using one-way ANOVA followed by Dunnett’s multiple comparisons test. Data are presented as mean ±SD from three independent biological replicates (n = 3). Statistical significance is denoted as ***p < 0.001, **p < 0.01, and *p < 0.05, compared to unstimulated control and ns as non-significant compared to unstimulated control.

## Discussion

The role of H3K4me3 in gene regulation remains unclear, with continued debate over whether it functions as an active regulator of gene expression or merely serves as a downstream marker of transcriptional activity. Although H3K4me3 is presumed to facilitate transcription initiation, evidence indicating that it is dispensable for transcription, complicates its functional interpretation^1–4^. In this study, we characterized the temporal dynamics of H3K4me3 during transcription initiation using the native, inducible human genes *TNF-α* and *IL-1β* as transcription models. By analysing gene activation at their endogenous loci, we preserved the native regulatory architecture and avoided potential artifacts commonly associated with transgene systems. Here, we show for the first time that H3K4me3 deposition at these promoters occurs predominantly as a post-transcriptional event, representing a downstream consequence of transcriptional activity rather than a prerequisite for initiation. To our knowledge, this is also a first demonstration of a histone modification being deposited at gene promoters after transcription initiation has already commenced. These findings advance our understanding of H3K4me3 function and call for a reassessment of its role in the regulation of transcriptional hierarchy.

Inducible gene models exhibit precise transcriptional on/off transitions, making them ideal for investigating associated histone modification dynamics. The rapid and transient expression kinetics of *TNF-α* and *IL-1β* genes^22–25^, coupled with short mRNA half-lives, enabled high-resolution analysis of early and delayed transcriptional effects, which are often obscured in constitutively expressed gene models. Our findings indicate that H3K4me3, a modification typically associated with active gene promoters, is not enriched at the *TNF-α* and *IL-1β* promoters during their peak transcriptional activity at 2 h time point, suggesting it is not required to initiate their transcription. Instead, we observed a delayed accumulation of H3K4me3, with its peak enrichment occurring around 6 h time point, after the transcriptional activity of *TNF-α* and *IL-1β* had subsided. This temporal uncoupling indicates a post-transcriptional deposition of H3K4me3 that is independent of active transcription at the time of its enrichment.

Further support for the post-transcriptional deposition of H3K4me3 was obtained through comparative analysis of additional histone modifications and transcriptional regulators. Histone acetylation marks are well established to closely associated with active transcription. We found that H3K27ac and H4K8ac, enrichment profiles were closely aligned with the transcription, validating the transcription profile and suggesting H3K4me3 is post-transcriptional downstream of H3K27ac. A previous study has demonstrated that targeted H3K27ac enrichment at promoters induced both H3K4me3 deposition and gene activation, whereas H3K4me3 recruitment failed to initiate transcription or promote H3K27ac. These findings are in line with current data showing that H3K4me3 is dispensable for transcription initiation and is deposited downstream of H3K27ac, consistent with a post-transcriptional role^35^. Additionally, phosphorylated RNA Polymerase II (Ser5 and Ser2), NF-κB, and p300 exhibited temporal patterns closely resembling transcriptional activity^20^, which markedly differed from the delayed accumulation of H3K4me3. Interestingly, the dynamic enrichment of MLL1, a methyltransferase responsible for H3K4me2 and H3K4me3^36^, shown to be involved in NF-κB mediated transcription^30–32^, closely aligned with the post-transcriptional enrichment profiles of H3K4me2 and H3K4me3 at *TNF-α* and *IL-1β* promoters. Although we found that H3K4me3 is deposited post-transcriptionally and independent of active transcription at the time of its writing, transcriptional inhibition significantly reduced its post-transcriptional accumulation, suggesting that H3K4me3 deposition still relies on prior transcriptional activity.

Our findings add to a growing body of evidence that challenges the canonical view of H3K4me3 as a requisite mark for transcription initiation. Despite its strong enrichment at active promoters, multiple studies indicate that H3K4me3 is dispensable for the onset of transcription. In mouse embryonic stem cells, deletion of CFP1 significantly reduces H3K4me3 at CpG island-associated promoters without affecting nascent or steady-state RNA levels^15,19^. Similarly, Drosophila cells lacking H3K4me3 maintain inducible gene expression, and yeast mutants devoid of H3K4 methylation remain viable and transcriptionally competent^16,18^. Transcriptional studies using reconstituted *in vitro* systems demonstrated that transcription is independent of enzymes responsible for catalysing H3K4me3^37^. Consistent with these findings, our inducible transcriptional model revealed that *MLL1* knockdown, which reduces H3K4me3 levels, does not impair *TNF-α* activation. These observations suggest that H3K4me3 is not essential for transcriptional initiation but may play a more nuanced role in transcriptional regulation post-transcriptionally. To our knowledge, this is the first study to offer a plausible explanation for the apparent disconnect between H3K4me3 enrichment at promoters and its non-essential role in transcription initiation. A recent study by Wang et al.^17^ supports this perspective, showing that depletion of H3K4me3 leads to only modest global changes in gene expression and is not required for RNA polymerase II loading at promoters. However, they observed increased RNA polymerase II pausing, suggesting that H3K4me3 regulates pause release. In contrast, our data show that H3K4me3 is deposited after the completion of the transcriptional cycle at inducible genes. When considered in the context of a constitutively active gene, this observation implies that H3K4me3 deposition may reflect regulation of subsequent transcriptional cycles. Together, these findings support a model in which H3K4me3 functions not as a universal driver of transcription initiation, but rather as a context-specific modulator of post-initiation events.

The observation of post-transcriptional H3K4me3 enrichment at inducible genes raises the possibility that a similar mechanism operates at constitutively expressed loci. However, dissecting this relationship is complicated by the continuous transcription and stable H3K4me3 levels at such genes, making it challenging to distinguish whether chromatin changes are a cause or a consequence of transcription, particularly when they occur after a temporal lag. To address this, we used a transcriptional termination model to test whether H3K4me3 is maintained at promoters following transcriptional shutdown. Using the constitutively expressed *MYC* gene, we confirmed transcriptional termination by a rapid decline in mRNA levels. Notably, H3K4me3 levels at the *MYC* promoter remained stable for at least four hours post-inhibition. This persistence suggests that H3K4me3 deposition is not strictly coupled to ongoing transcription but may instead reflect a delayed, post-initiation event dependent on prior transcriptional activity. For instance, genes marked by H3K4me3 but lacking active transcription likely reflect prior transcriptional activity^38^. These findings are consistent with previous reports of sustained H3K4me3 after transcriptional arrest, suggesting that its post-transcriptional retention may reflect a broader regulatory mechanism^39^.

Although our study focuses on a limited number of genes, it uncovers a previously underappreciated role for H3K4me3 in post-initiation transcriptional regulation. To evaluate how widespread this mechanism, future studies will need to investigate the dynamics and regulatory roles of H3K4me3 across a broader range of active promoters, ideally at genome-wide resolution. However, based on the current literature, we propose that H3K4me3 may serve several critical roles following transcription initiation. These include: (i) stabilizing promoter-proximal open chromatin and acting as a barrier element to prevent the encroachment of repressive or silencing histone modifications^40,41^; (ii) facilitating transcriptional elongation and modulating RNA turnover through interactions with the transcriptional machinery at select constitutively active genes^17,20,42^; (iii) contributing to genome integrity^43,44^; and (iv) supporting transcriptional memory^39^.

In conclusion, our findings establish H3K4me3 as a transcription-dependent, post-transcriptional epigenetic mark. By positioning it as a consequence rather than a prerequisite of transcriptional activity, we redefine the current understanding of its regulatory function. This study opens new avenues for elucidating the role of H3K4me3 in shaping chromatin architecture and maintaining transcriptional fidelity, thereby prompting a critical re-evaluation of long-standing paradigms. These insights may have important implications for interpreting the role of H3K4me3 in diseases, particularly in pathological contexts marked by its aberrant modification, and for developing therapeutic strategies targeting epigenetic dysregulation.

## Methods

### Cell culture

THP-1 monocytic cells (ATCC) were cultured in suspension in RPMI-1640 medium (Gibco, Cat# 23-400-021) supplemented with 10% heat-inactivated fetal bovine serum (FBS) (Gibco, Cat# 10270-106) and 1% of penicillin–streptomycin (Gibco, Cat# 15140-122). Cells were maintained in a humidified incubator at 37 °C with 5% CO2 and sub-cultured every 2–3 days. For transcriptional activation studies, THP-1 monocytes (1×10^7^ cells) were cultured in 10 mL of complete RPMI medium and induced with *E. coli* lipopolysaccharide (LPS; 500 ng/mL) (Sigma Aldrich, Cat# L4005) for various durations (0, 0.5, 2, 6, 12, and 24 h). At each time point, 1 mL of the culture was harvested for RNA isolation, while the remaining 9 mL were processed for chromatin immunoprecipitation (ChIP) assays. For lentiviral production, HEK293T cells were cultured in Dulbecco’s Modified Eagle Medium (DMEM; Gibco, Cat# 12100-061) supplemented with 10% FBS and 1% penicillin–streptomycin solution. Cells were cultured under the same conditions as THP-1 cells.

### Lentivirus plasmid construction and transfection

shRNA sequences targeting human *MLL1* were designed based on the Broad Institute’s TRC (Transgenic RNAi Consortium) shRNA design process^45^ and oligonucleotides were synthesized and annealed (Supplementary table 2). The resulting duplexes were ligated into the pLKO.1-vector (Addgene# 8453), linearized with *Age* I and *EcoR* I, using Quick T4 DNA Ligase (NEB, Cat# M2200L) kit. HEK293T cells were transfected with either sh*MLL1* or an empty vector (control) for lentiviral production along with third-generation lenti-packaging plasmids pREV, pRRE, and pMD2.G using Xfect™ Transfection Reagent (Takara, Cat# 631318) based on the manufacturer’s protocol. After 48 h and 72 h of post-transfection, viral particles were collected. Lentiviral transduction was performed on THP-1 cells (5 × 10^5^ cells/mL) in the presence of 8 μg/mL polybrene (Sigma-Aldrich, Cat# 107689). After 24 h, cells were added with fresh RPMI-1640 medium, and puromycin (Sigma-Aldrich, Cat# P4512) selection was carried out with 1 μg/mL concentration for 7 days. Later MLL1 knockdown efficiencies were evaluated by western blot analysis and subsequently these cells were used to analyse LPS-induced activation of *TNF-α* transcription.

### RNA extraction, cDNA synthesis and Real-time quantitative PCR (qRT-PCR)

Total RNA was isolated using RNAiso Plus (Takara, Cat# 9109) according to the manufacturer’s instructions. PrimeScript RT reagent kit (Takara, Cat# 6110A) was used to prepare cDNA from 1 μg of RNA. Reverse transcription quantitative PCR (RT–qPCR) was performed with Takara SYBR Green Master Mix (Cat# RR820) using gene-specific primers (listed in Supplementary Table 1) on an Applied Biosystems 7900HT Real-Time PCR System. mRNA expression levels were normalized to *β2-microglobulin* (*β2M*) as a housekeeping control, and relative quantification was calculated using a ΔΔ*C*_t_ method. Statistical analysis was performed by GraphPad Prism v.9 software (GraphPad).

### Chromatin immunoprecipitation (ChIP)

ChIP assays were performed as previously described^46^. Briefly, THP-1 cells were fixed with 1% formaldehyde for 10 min at room temperature, and crosslinking was quenched with 0.125 M glycine. Chromatin was prepared by lysing the cells in ChIP lysis buffer (20 mM HEPES, pH 7.6, 1% SDS, 1 mM sodium butyrate, protease inhibitors) on ice. Chromatin was sonicated using a bath sonicator (SONOREX) to yield DNA fragments of approximately 200–600 bp. The lysate was pre-cleared by centrifugation, and 10% of the input was reserved for normalization. For each ChIP reaction, 20 µl of Protein A Dynabeads (Invitrogen, Cat# 10002D) were coated with 1.5 µg of the respective antibody (listed in Supplementary Table 2), diluted in 0.1% BSA-containing ChIP dilution buffer (16.7 mM Tris-HCl (pH 8.0), 1.2 mM EDTA, 167 mM NaCl, 1% Triton X-100), and incubated for 2 h at room temperature with gentle rotation. The antibody-coated beads were then added to the pre-cleared chromatin (equivalent to 1 × 10^6^ cells per sample) in ChIP dilution buffer containing 0.01% SDS and incubated at 4 °C overnight with rotation. Next day, magnetic beads containing the immunoprecipitated chromatin complexes were separated using a magnetic stand. Beads were then sequentially washed with ice-cold buffers: once with low-salt wash buffer (20 mM Tris-HCl, pH 8.0, 2 mM EDTA, 150 mM NaCl, 0.1% SDS, 1% Triton X-100), twice with high-salt wash buffer (20 mM Tris-HCl, pH 8.0, 2 mM EDTA, 500 mM NaCl, 0.1% SDS, 1% Triton X-100), and twice with TE buffer (10 mM Tris-HCl, pH 8.0, 1 mM EDTA). Chromatin was eluted in elution buffer (10 mM Tris-HCl, pH 8.0, 5 mM EDTA, 300 mM NaCl, 0.5% SDS) supplemented with proteinase K (20 mg/ml) and incubated overnight at 65 °C for de-crosslinking the chromatin. ChIP DNA was purified using the PCR Clean-up/Gel Extraction Kit (Genetix, Cat# NP-36107). Enrichment of target genomic regions was analysed by real-time PCR using promoter-specific primers listed in Supplementary Table 1. Data were normalized to input DNA and quantified using the ΔCt method.

### Transcription inhibition analysis using Actinomycin D

To investigate the transcription dependence of chromatin modifications, THP-1 cells (1 × 10^6^ cells per time point) were pre-treated with 1 μM Actinomycin D (Sigma-Aldrich, Cat# A4262) for 30 min to inhibit RNA synthesis. Following pre-treatment, cells were induced with LPS, (500 ng/mL) for 0, 2, and 6 h. and harvested for downstream analysis of *TNF-α* and *IL-1β* mRNA expression by qRT-PCR and for H3K4me3 ChIP enrichment at the *TNF-α* and *IL-1β* promoter, as described. Untreated cells (no ActD) subjected to identical LPS stimulation served as experimental controls.

To evaluate whether the post-transcriptional accumulation of H3K4me3 represents a general chromatin regulatory mechanism (Fig. 5), THP-1 cells (1 × 10^6^ per ml) were treated with 1 μM Actinomycin D (Sigma-Aldrich, Cat# A4262) for varying durations (0, 1, 2, 6, and 8 h). Cells were harvested at each time point for downstream analysis of *TNF-α* mRNA expression by qRT-PCR and for H3K4me3 chromatin immunoprecipitation (ChIP) enrichment at the *TNF-α* promoter. Untreated cells (0 h) served as the control condition.

### Western Blot

Total proteins were isolated by lysing cells in RIPA buffer (Clontech, Cat# ST0349) supplemented with Halt™ Protease Inhibitor Cocktail (Takara, Cat# 635673) and Phosphatase Inhibitor Cocktail (Sigma-Aldrich, Cat# P0044). Protein concentration was estimated using the BCA Protein Assay Kit (Thermo Scientific, Cat# 71285-3). Equal amounts of protein (30 µg per sample) were resolved by SDS–PAGE on 15% and 8% acrylamide gels (Bio-Rad) and transferred onto nitrocellulose membranes (Amersham™ Hybond™, Cat# 10600023). Membranes after protein transfer blocked with 5% BSA with 1× TBST (137 mM NaCl, 2.7 mM KCl, 20 mM Tris-HCl (pH 7.4), 0.1% Tween-20), for 1 h at room temperature and later incubated at 4 °C overnight with the respective primary antibodies against GAPDH, MLL1, and H3K4me3 (Cat. No. listed in Supplementary Table 2). After washing with 1× TBST, membranes were incubated with HRP-conjugated goat anti-rabbit IgG (Jackson Immuno Research Laboratories Cat# AB_2313567) secondary antibodies. Signal was developed using the ECL Select™ Western Blotting Detection Kit (Amersham™, Cat# RPN2235) and visualized with the ChemiDoc Imaging System (Bio-Rad). Relative band intensities were quantified using ImageJ software. Original blots are provided in Supplementary Fig. 5.

### Statistical Significance

Statistical analysis for qRT–PCR and ChIP enrichment assays was performed using one-way ANOVA with Dunnett’s multiple comparisons test using GraphPad Prism v9.0. Data are presented as mean ± SD and SEM from three independent biological replicates. Statistical significance is denoted as *p < 0.05, **p < 0.001, and ***p < 0.0001, compared to control cells, ###p < 0.001, ##p < 0.01, and #p < 0.05 compared to control LPS 2 h stimulated cells, $$$P < 0.001, $$P < 0.01, $P < 0.05 compared to LPS 6 h stimulated cells and ns as non-significant compared to unstimulated (control) cells. Statistical methods and data analysis are detailed in the figure legends. Analyses of western blot in Fig 5a was independently performed three times with similar findings.

## Abbreviations

H3K4me3: Histone H3 Lysine 4 trimethylation
H3K4me2: Histone H3 Lysine 4 dimethylation
H3K4me1: Histone H3 Lysine 4 monomethylation
H3K27ac: Histone H3 Lysine 27 Acetylation
TNF-α: Tumor Necrosis Factor Alpha
IL-1β: Interleukin 1 Beta
MLL1: Mixed Lineage Leukemia 1
Pol II: RNA Polymerase II
NF-κB: Nuclear Factor kappa-light-chain-enhancer of activated B cells
p300: Histone Acetyltransferase p300
ActD: Actinomycin D
MYC: MYC Proto-Oncogene
LPS: Lipopolysaccharide
ChIP: Chromatin Immunoprecipitation
TSS: Transcription Start Site
qRT-PCR: Quantitative Reverse Transcription PCR
h: Hours

## Acknowledgements

The authors thank the Director, CSIR-IICT, Dr. Sistla Ramakrishna, Dr. Shasi Vardhan Kalivendi, HOD, Applied Biology, CSIR-IICT, Hyderabad, India for providing the facilities necessary for the conducting of this research work. CSIR-IICT manuscript communication number: IICT/Pubs./2025/207. K.P.W acknowledges DST Inspire, Govt. of India for providing SRF. S.C. acknowledges the DBT-Ramalingaswami Re-entry Fellowship, Govt. of India, New Delhi. We thank Dr. Sai Balaji Andugulapati, IICT for support.

## Author contributions

Conceptualization: S.C; Methodology: K.P.W, S.C; *in vitro* cell culture: K.P.W, S.C; Manuscript writing - review & editing: S.C; Funding acquisition: S.C.

## Data availability statement

The data that support the findings of this study are available from the corresponding author upon reasonable request.

## References

1 Haoming, Y. & Bluma, J. L. a. Functional Roles of H3K4 Methylation in Transcriptional Regulation. Molecular and cellular biology 44, 505--515, doi:10.1080/10985549.2024.2388254 (2024).

2 Howe, F. S., Fischl, H., Murray, S. C. & Mellor, J. Is H3K4me3 instructive for transcription activation? BioEssays : news and reviews in molecular, cellular and developmental biology 39, 1–12, doi:10.1002/bies.201600095 (2017).

3 Hua, W. & Kristian, H. Roles of H3K4 methylation in biology and disease. Trends in cell biology 35, 115–128, doi:10.1016/j.tcb.2024.06.001 (2025).

4 Talbert, P. B. & Henikoff, S. The Yin and Yang of Histone Marks in Transcription. Annual review of genomics and human genetics 22, 147–170, doi:10.1146/annurev-genom-120220-085159 (2021).

5 Millán-Zambrano, G., Burton, A., Bannister, A. J. & Schneider, R. Histone post-translational modifications - cause and consequence of genome function. 23, 563–580, doi:10.1038/s41576-022-00468-7 (2022).

6 Weinzapfel, E. N. & Fedder-Semmes, K. N. Beyond the tail: the consequence of context in histone post-translational modification and chromatin research. 481, 219–244, doi:10.1042/bcj20230342 (2024).

7 Zhou, V. W., Goren, A. & Bernstein, B. E. Charting histone modifications and the functional organization of mammalian genomes. Nature reviews. Genetics 12, 7–18, doi:10.1038/nrg2905 (2011).

8 Allis, C. D. & Jenuwein, T. The molecular hallmarks of epigenetic control. Nature reviews. Genetics 17, 487–500, doi:10.1038/nrg.2016.59 (2016).

9 Bernstein, B. E. et al. Genomic maps and comparative analysis of histone modifications in human and mouse. Cell 120, 169–181, doi:10.1016/j.cell.2005.01.001 (2005).

10 Pokholok, D. K. et al. Genome-wide map of nucleosome acetylation and methylation in yeast. Cell 122, 517–527, doi:10.1016/j.cell.2005.06.026 (2005).

11 Santos-Rosa, H. et al. Active genes are tri-methylated at K4 of histone H3. Nature 419, 407–411, doi:10.1038/nature01080 (2002).

12 Schneider, R. et al. Histone H3 lysine 4 methylation patterns in higher eukaryotic genes. Nature cell biology 6, 73–77, doi:10.1038/ncb1076 (2004).

13 Lauberth, S. M. et al. H3K4me3 interactions with TAF3 regulate preinitiation complex assembly and selective gene activation. Cell 152, 1021–1036, doi:10.1016/j.cell.2013.01.052 (2013).

14 Vermeulen, M. et al. Selective anchoring of TFIID to nucleosomes by trimethylation of histone H3 lysine 4. Cell 131, 58–69, doi:10.1016/j.cell.2007.08.016 (2007).

15 Clouaire, T. et al. Cfp1 integrates both CpG content and gene activity for accurate H3K4me3 deposition in embryonic stem cells. Genes & development 26, 1714–1728, doi:10.1101/gad.194209.112 (2012).

16 Luis, M. S. & Stephen, B. Yeast Swd2 Is Essential Because of Antagonism between Set1 Histone Methyltransferase Complex and APT (Associated with Pta1) Termination Factor*. Journal of Biological Chemistry 287, 15219–15231, doi:10.1074/jbc.M112.341412 (2012).

17 Wang, H. et al. H3K4me3 regulates RNA polymerase II promoter-proximal pause-release. Nature 615, 339–348, doi:10.1038/s41586-023-05780-8 (2023).

18 Hödl, M. & Basler, K. Transcription in the absence of histone H3.2 and H3K4 methylation. Current biology : CB 22, 2253–2257, doi:10.1016/j.cub.2012.10.008 (2012).

19 Carlone, D. L. et al. Reduced genomic cytosine methylation and defective cellular differentiation in embryonic stem cells lacking CpG binding protein. Molecular and cellular biology 25, 4881–4891, doi:10.1128/mcb.25.12.4881-4891.2005 (2005).

20 Ding, Y. et al. ATX1-generated H3K4me3 is required for efficient elongation of transcription, not initiation, at ATX1-regulated genes. PLoS genetics 8, e1003111, doi:10.1371/journal.pgen.1003111 (2012).

21 Henikoff, S. & Shilatifard, A. Histone modification: cause or cog? Trends in genetics : TIG 27, 389–396, doi:10.1016/j.tig.2011.06.006 (2011).

22 Adamik, J. et al. Distinct mechanisms for induction and tolerance regulate the immediate early genes encoding interleukin 1β and tumor necrosis factor α. PloS one 8, e70622, doi:10.1371/journal.pone.0070622 (2013).

23 Pulugulla, S. H., Adamik, J., Grillini, A. N., Galson, D. L. & Auron, P. E. Specific transcription factors distinctly regulate kinetics of IL1B and TNF gene expression. The Journal of Immunology 196, 189.114–189.114, doi:10.4049/jimmunol.196.Supp.189.14 (2016).

24 Sharif, O., Bolshakov, V. N., Raines, S., Newham, P. & Perkins, N. D. Transcriptional profiling of the LPS induced NF-κB response in macrophages. BMC Immunology 8, 1, doi:10.1186/1471-2172-8-1 (2007).

25 Suzuki, T. et al. Comprehensive gene expression profile of LPS-stimulated human monocytes by SAGE. Blood 96, 2584–2591, doi:10.1182/blood.V96.7.2584 (2000).

26 Stasevich, T. J. et al. Regulation of RNA polymerase II activation by histone acetylation in single living cells. Nature 516, 272–275, doi:10.1038/nature13714 (2014).

27 Wang, Z. et al. Combinatorial patterns of histone acetylations and methylations in the human genome. Nature genetics 40, 897–903, doi:10.1038/ng.154 (2008).

28 Hsin, J. P. & Manley, J. L. The RNA polymerase II CTD coordinates transcription and RNA processing. Genes & development 26, 2119–2137, doi:10.1101/gad.200303.112 (2012).

29 Bhatt, D. & Ghosh, S. Regulation of the NF-κB-Mediated Transcription of Inflammatory Genes. Frontiers in immunology 5, 71, doi:10.3389/fimmu.2014.00071 (2014).

30 Kimball, A. S. et al. The Histone Methyltransferase MLL1 Directs Macrophage-Mediated Inflammation in Wound Healing and Is Altered in a Murine Model of Obesity and Type 2 Diabetes. 66, 2459–2471, doi:10.2337/db17-0194 (2017).

31 Wang, X. et al. MLL1, a H3K4 methyltransferase, regulates the TNFα-stimulated activation of genes downstream of NF-κB. Journal of cell science 125, 4058–4066, doi:10.1242/jcs.103531 (2012).

32 Wang, J. et al. Histone modifications and their roles in macrophage-mediated inflammation: a new target for diabetic wound healing. Frontiers in immunology 15, 1450440, doi:10.3389/fimmu.2024.1450440 (2024).

33 Koba, M. & Konopa, J. [Actinomycin D and its mechanisms of action]. Postepy higieny i medycyny doswiadczalnej (Online) 59, 290–298 (2005).

34 Herrick, D. J. & Ross, J. The half-life of c-myc mRNA in growing and serum-stimulated cells: influence of the coding and 3’ untranslated regions and role of ribosome translocation. Molecular and cellular biology 14, 2119–2128, doi:10.1128/mcb.14.3.2119-2128.1994 (1994).

35 Zhao, W. et al. Investigating crosstalk between H3K27 acetylation and H3K4 trimethylation in CRISPR/dCas-based epigenome editing and gene activation. Scientific Reports 11, 15912, doi:10.1038/s41598-021-95398-5 (2021).

36 Dou, Y. et al. Regulation of MLL1 H3K4 methyltransferase activity by its core components. Nature structural & molecular biology 13, 713–719, doi:10.1038/nsmb1128 (2006).

37 Pavri, R. et al. Histone H2B monoubiquitination functions cooperatively with FACT to regulate elongation by RNA polymerase II. Cell 125, 703–717, doi:10.1016/j.cell.2006.04.029 (2006).

38 Guenther, M. G., Levine, S. S., Boyer, L. A., Jaenisch, R. & Young, R. A. A chromatin landmark and transcription initiation at most promoters in human cells. Cell 130, 77–88, doi:10.1016/j.cell.2007.05.042 (2007).

39 Ng, H. H., Robert, F., Young, R. A. & Struhl, K. Targeted recruitment of Set1 histone methylase by elongating Pol II provides a localized mark and memory of recent transcriptional activity. Molecular cell 11, 709–719, doi:10.1016/s1097-2765(03)00092-3 (2003).

40 Ma, M. K., Heath, C., Hair, A. & West, A. G. Histone crosstalk directed by H2B ubiquitination is required for chromatin boundary integrity. PLoS genetics 7, e1002175, doi:10.1371/journal.pgen.1002175 (2011).

41 Wysocka, J. et al. A PHD finger of NURF couples histone H3 lysine 4 trimethylation with chromatin remodelling. Nature 442, 86–90, doi:10.1038/nature04815 (2006).

42 Hu, S. et al. H3K4me2/3 modulate the stability of RNA polymerase II pausing. 33, 403–406, doi:10.1038/s41422-023-00794-3 (2023).

43 Abu-Zhayia, E. R. et al. CDYL1-dependent decrease in lysine crotonylation at DNA double-strand break sites functionally uncouples transcriptional silencing and repair. Molecular cell 82, 1940–1955.e1947, doi:10.1016/j.molcel.2022.03.031 (2022).

44 Gong, F., Clouaire, T. & Aguirrebengoa, M. Histone demethylase KDM5A regulates the ZMYND8-NuRD chromatin remodeler to promote DNA repair. 216, 1959–1974, doi:10.1083/jcb.201611135 (2017).

45 Moffat, J. et al. A lentiviral RNAi library for human and mouse genes applied to an arrayed viral high-content screen. Cell 124, 1283–1298, doi:10.1016/j.cell.2006.01.040 (2006).

46 Ansalone, C. et al. TNF is a homoeostatic regulator of distinct epigenetically primed human osteoclast precursors. 80, 748–757, doi:10.1136/annrheumdis-2020-219262 (2021).

