## Supplementary figures and tables for "H3K4me3 is a post-transcriptional histone mark"

#### Supplementary Figure 1

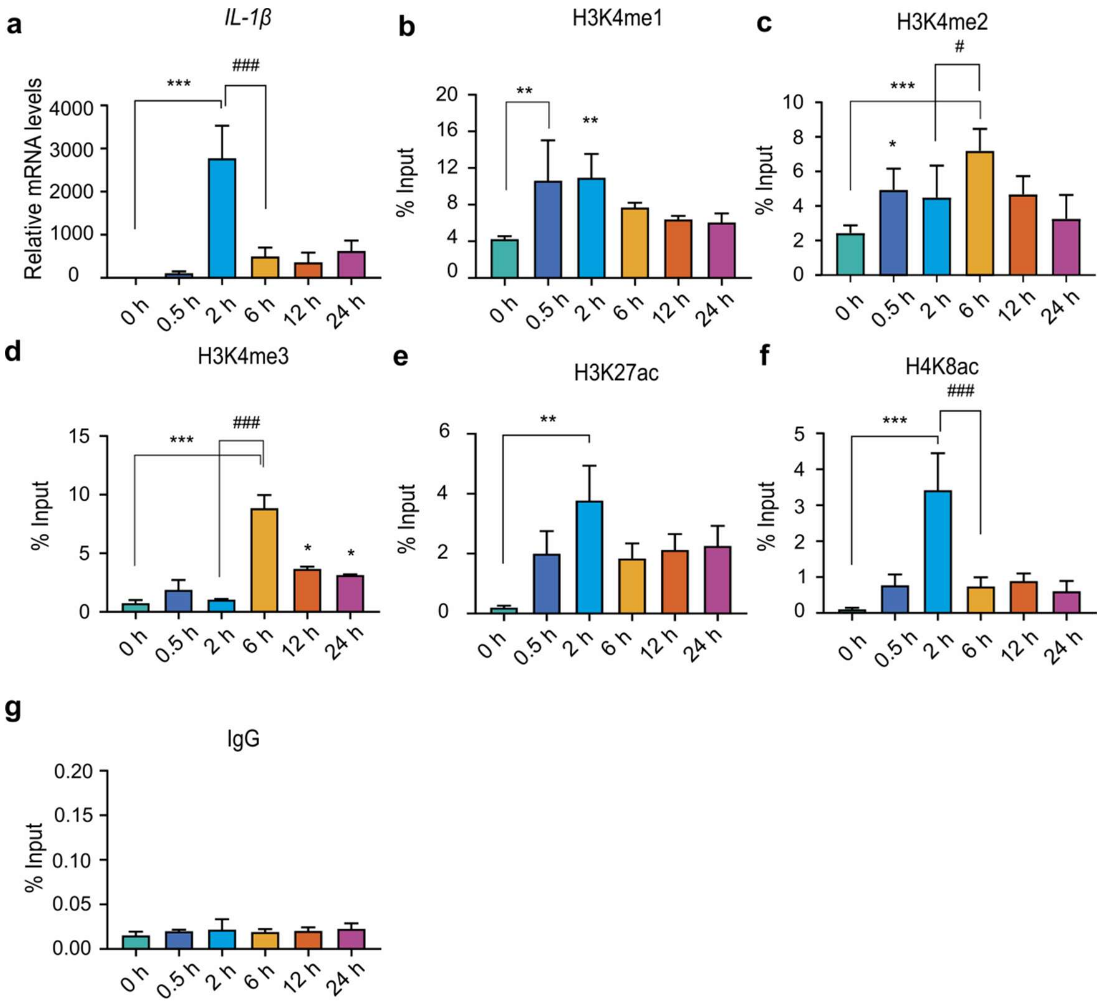


**Supplementary Figure 1. Inducible gene activation model reveals temporal decoupling of H3K4me3 deposition and transcription.** (**a-g**) Cells were stimulated with LPS (500 ng/ml) and harvested at 0, 0.5, 2, 6, 12, and 24 h post-stimulation for transcriptional and chromatin immunoprecipitation (ChIP) analyses. (**a**), qPCR analysis of *IL-1β* mRNA levels following LPS stimulation, used as readout of transcriptional activation. (**b-g**), ChIP enrichment analysis was performed using antibodies against (**b**) H3K4me1, (**c**) H3K4me2, (**d**) H3K4me3, (**e**) H3K27ac, (**f**) H4K8ac, and (**g**) IgG control. Immunoprecipitated DNA was analysed by qPCR using primers specific to the *IL-1β* promoter. Statistical analysis was performed using one-way ANOVA followed by Dunnett's multiple comparisons test. Data are presented as mean ± SD (n=3 for a, b, c and f) and SEM (n = 4 for e) independent biological replicates. Statistical significance is indicated as ***$p < 0.001$, **$p < 0.01$, and *$p < 0.05$, relative to the unstimulated control and ###P < 0.001, ##P < 0.01, #P < 0.05 compared to LPS 2 h stimulated cells.

#### Supplementary Figure 2

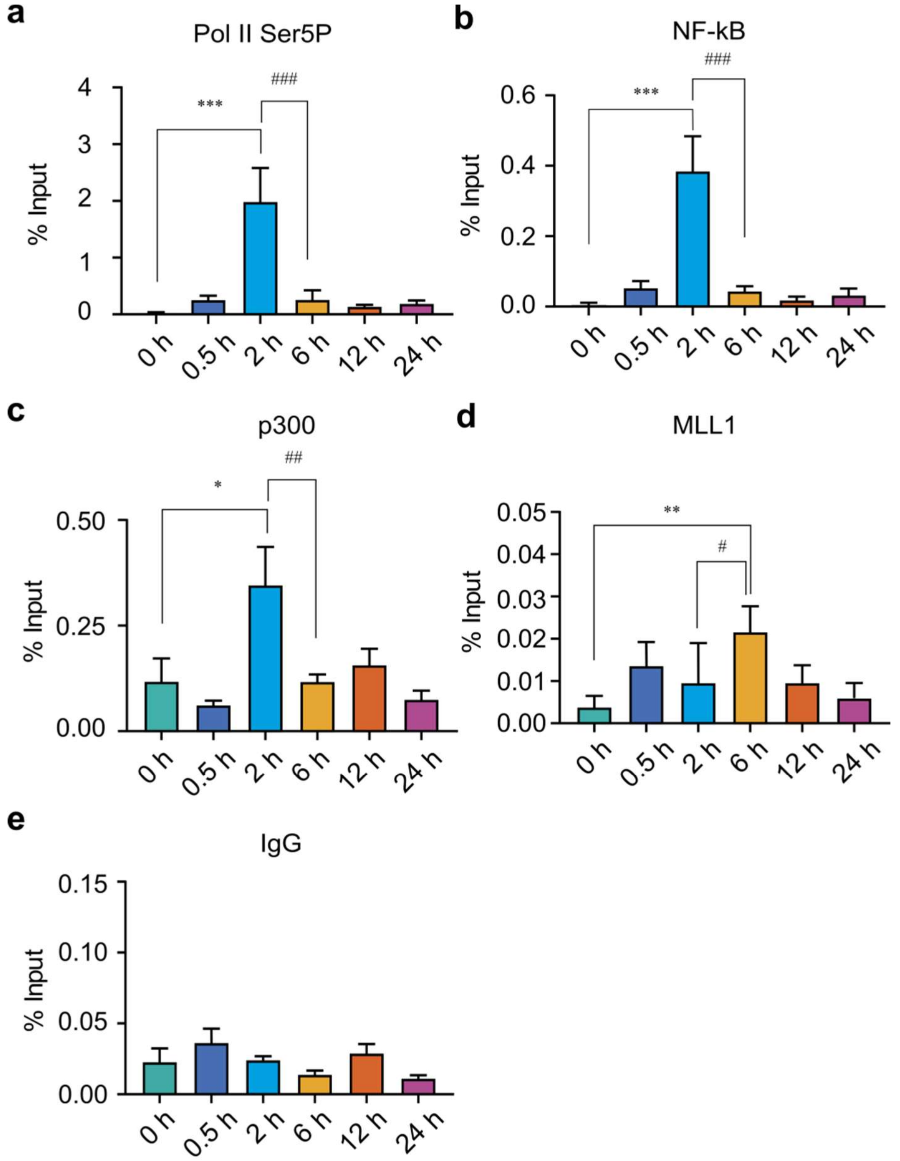


**Supplementary Figure 2. ChIP enrichment analysis of transcriptional regulators reveals temporal divergence from H3K4me3 dynamics during LPS-induced gene activation.** (**a–f**), THP-1 cells were stimulated with LPS (500 ng/ml) and harvested at 0, 0.5, 2, 6, 12, and 24 h to assess temporal occupancy of key transcriptional regulators at the *IL-1β* locus. ChIP enrichment analysis was performed using antibodies against (a) RNA polymerase II Ser5-phosphorylated (Pol II Ser5P), marking transcriptional initiation. (b) NF-κB (p65), a stimulus-responsive transcription factor; (c) MLL1, a histone methyltransferase responsible for H3K4me3 methylation; (d) P300, a histone acetyltransferase linked to transcriptional coactivation; and (e) IgG as a negative control. Enriched chromatin was quantified by qPCR using primers specific to the *IL-1β* promoter. Statistical analysis was performed using one-way ANOVA followed by

Dunnett's multiple comparisons test. Data are presented as mean ± SD (n=3 for b and e), SEM (n=4 for a, c and n=3 for d) independent biological replicates. Statistical significance is indicated as ***$p < 0.001$, **$p < 0.01$, and *$p < 0.05$, compared to unstimulated control

#### Supplementary Figure 3

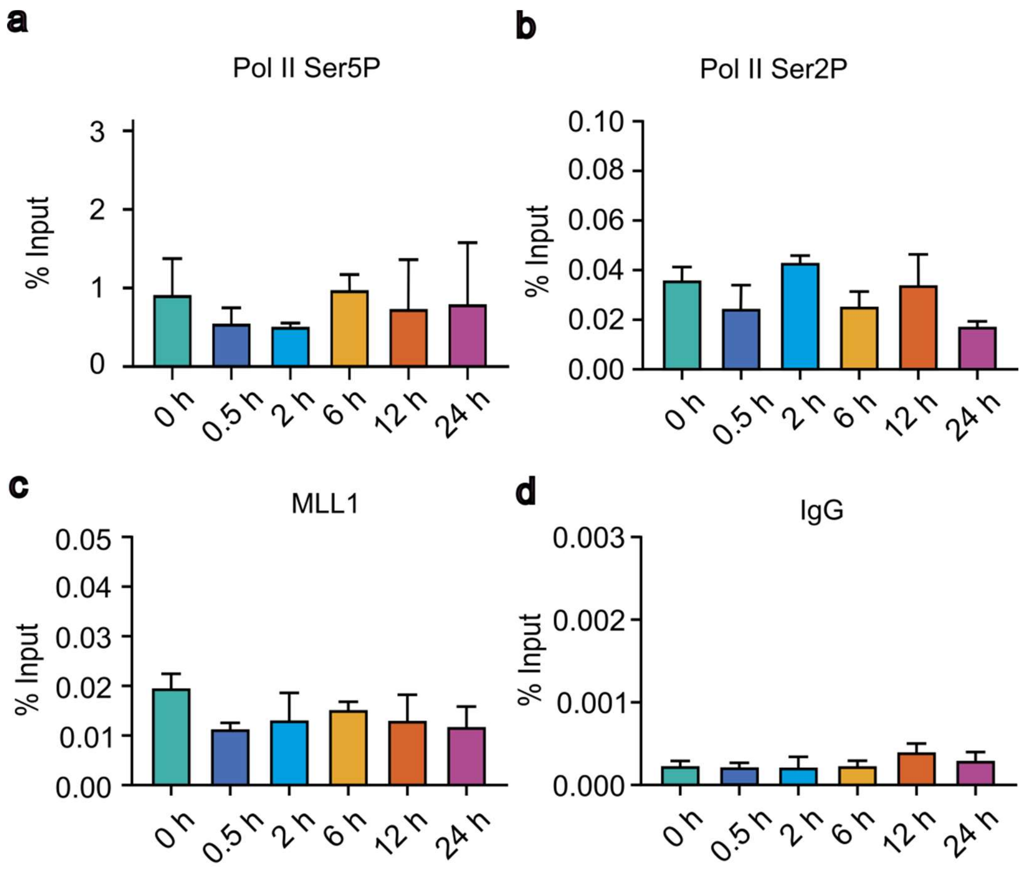


**Supplementary Figure 3. Validation of ChIP assay specificity and transcriptional model in THP-1 cells.**THP-1 cells were treated with LPS for indicated time points 0, 0.5, 2, 6, 12, and 24 h and ChIP enrichment assays were performed using antibodies against (a) RNA polymerase II Ser5-phosphorylated (Pol II Ser5P), (b) RNA polymerase II Ser2-phosphorylated (Pol II Ser2P), (c) MLL1; and (d) IgG (negative control). Immunoprecipitated DNA was analysed by qPCR using primers specific to the *GAPDH* promoter. Statistical analysis was performed using one-way ANOVA followed by Dunnett's multiple comparisons test. Data are presented as mean ± SD (n=3) three independent biological replicates.

### Supplementary Figure 4

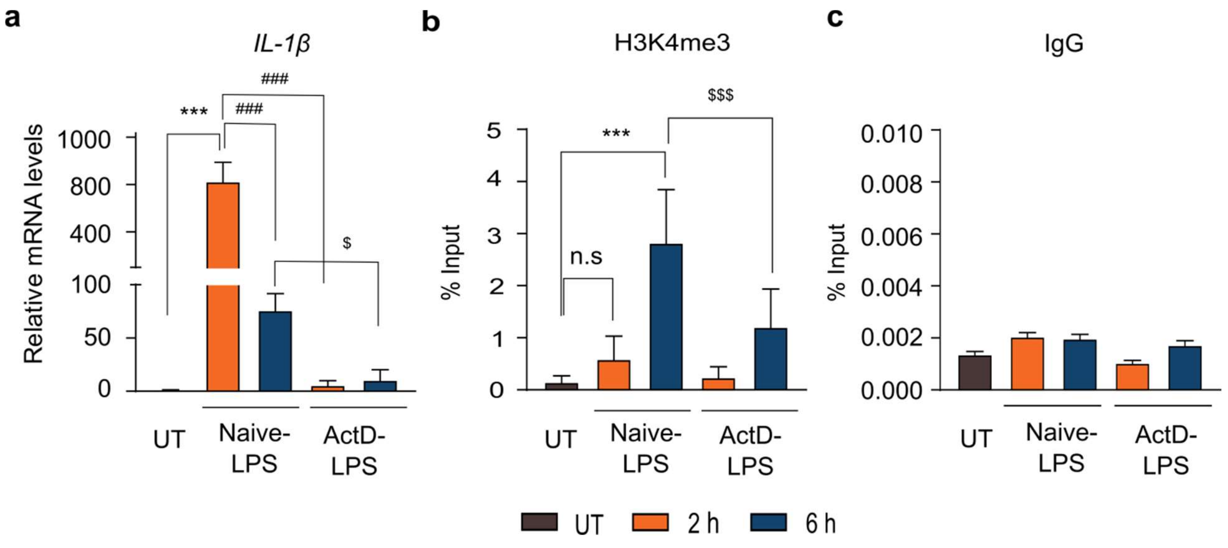


**Supplementary Figure 4. Post-transcriptional H3K4me3 loading requires active transcription.** To determine whether active transcription is necessary for the post-transcriptional deposition of H3K4me3 at gene promoter, THP-1 monocytic cells were pre-treated with the transcriptional inhibitor Actinomycin D (1 µM) for 30 min prior to stimulation with LPS (500 ng/mL), and cells were harvested at 0 h, 2 h, and 6 h post-stimulation for analysis of gene expression and histone modifications. (**a**) qPCR analysis of *IL-1β* mRNA levels in control and Actinomycin D-treated THP-1 cells following LPS stimulation. (**b**) ChIP analysis using antibodies against H3K4me3 to assess enrichment at the *IL-1β* promoter. (**c**) ChIP with normal IgG was included as a negative control. Statistical analysis was performed using one-way ANOVA followed by Dunnett's multiple comparisons test. Data are presented as mean ± SD. from three biological replicates (n = 3). Statistical significance is denoted as P < 0.001, P < 0.01, P < 0.05 versus unstimulated control, ###P < 0.001, ##P < 0.01, #P < 0.05 compared to LPS 2 h stimulated cells, $$$P < 0.001, $$P < 0.01, $P < 0.05 compared to LPS 6 h stimulated cells and ns as non-significant compared to unstimulated control.

### Supplementary Figure 5

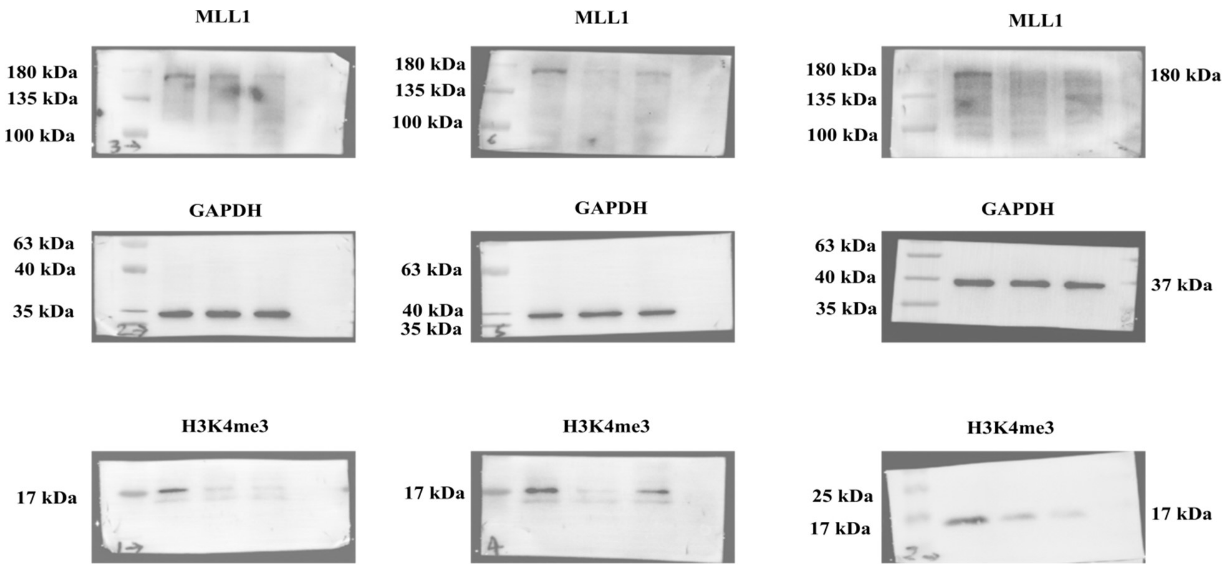


**Supplementary Figure 5:** Uncropped Western blots images showing protein levels of MLL1, H3K4me3, and GAPDH in THP-1 cells transduced with shMLL1 (n=2). Western was performed independently three times (n=3). MLL1 knockdown efficiency was validated by reduced MLL1 protein levels and corresponding decrease in H3K4me3, a downstream histone methylation mark. GAPDH was used as a loading control. These blots correspond to data presented in 5a.

**Supplementary Table 1: Oligonucleotide sequences used in ChIP-qPCR and RT-qPCR analysis.**

| Gene | Forward primer sequence | Reverse primer sequence |
|---|---|---|
| *β2M* | 5'- AGGCTATCCAGCGTACTCCA-3' | 5- 'CGGATGGATGAAACCCAGACA-3’ |
| *TNF-α* | 5'- CTCTTCTGCCTGCTGCACTTTG-3' | 5'- ATGGGCTACAGGCTTGTCACTC-3' |
| *IL-1β* | 5'- CCACAGACCTTCCAGGAGAA-3' | 5'- GTGCAGTTCAGTGGATCGTACAGG-3' |
| *MYC* | 5'- CCTGGTGCTCCATGAGGAGAC-3' | 5'- CAGACTCTGACCTTTTGCCAGG-3' |
| *TNF-α* promoter | 5'- ATCAGTCAGTGGCCCAGAAG-3' | 5'- TCATCTGGAGGAAGCGGTAG-3' |
| *IL-1β* promoter | 5'- ACCTTGGGTGCTGTTCTCTG-3' | 5'- CTGGTCTTGCAGGGTTGTGT-3' |
| *TNF-α* Exon 4 | 5'-TCTCCTACCAGACCAAGGTC -3' | 5'- CAAAGTAGACCTGCCCAGGAC-3' |
| *GAPDH* Promoter | 5'- TCATCCAAGCGTGTAAGGGT-3' | 5'- ACTGAGATTGGCCCGATGG-3' |
| *MYC* promoter | 5'- GGGACTTCTTGATCAAAGCGC-3' | 5'- CGCATCCTTGTCCTGTGAGT-3' |

**Supplementary Table 2: shRNA oligonucleotide sequences used for *MLL1* knockdown.**

| | | |
|---|---|---|
| *ShMLL1# 1* | 5'- CCGGGAGCTGTAAACATTCACATCTTCG CAGAAGATGTAGGATTTAACAGTGCTTTTT G-3’ | 5'- AATTCAAAAAGCACTGTTAAATCCTACATC TTCTGCGAAGATGTGAATGTTTACAGCTG CC-3' |
| *ShMLL1# 2* | 5'- CCGGCGCGCGTGATTACTCAATTTAACTCG GATAATAATGGAATACGGAATACCGGCTTT TG-3' | 5'- AATTCAAAAAGCCGGTATTTCCGTATTCCA TTATTATCCGAGTTAAATTGAGTAATCACG CGCG-3' |

**Table S3: List of antibodies used in this study.**

| S. No | Antibody | Catalogue No |
|---|---|---|
| 1 | GAPDH (CST) | 5174 |
| 2 | H3K4me1 (Puregene) | 73303 |
| 3 | H3K4me2 (Millipore) | 07-030 |
| 4 | H3K4me3 (CST) | 9751 |
| 5 | H3K27ac (CST) | 8173 |
| 6 | H4K8ac (Millipore) | 07-328 |
| 7 | Phospho-Rpb1 CTD (Ser2) (CST) | 13499 |
| 8 | Anti-RNA polymerase II CTD repeat YSPTSPS (phospho S5) (Abcam) | Ab5408 |
| 9 | NF-κB (CST) | 3033 |
| 10 | MLL-1 (CST) | 34907 |
| 11 | P300 (CST) | 54062 |
| 12 | IgG (CST) | 2729 |
